# Fluorescence-based 3D targeting of FIB-SEM acquisition of small volumes in large samples

**DOI:** 10.1101/2021.01.18.427072

**Authors:** Paolo Ronchi, Pedro Machado, Edoardo D’Imprima, Giulia Mizzon, Benedikt T. Best, Lucia Cassella, Sebastian Schnorrenberg, Marta G. Montero, Martin Jechlinger, Anne Ephrussi, Maria Leptin, Julia Mahamid, Yannick Schwab

## Abstract

Cells are three dimensional objects. Therefore, 3D electron microscopy is often crucial for correct interpretation of ultrastructural data. Today samples are frequently imaged in 3D at ultrastructural resolution using volume Scanning Electron Microscopy (SEM) methods such as Focused Ion Beam (FIB) SEM and Serial Block face SEM. While these imaging modalities allow for automated data acquisition, precise targeting of (small) volumes of interest within a large sample remains challenging. Here, we provide an easy and reliable approach to target FIB-SEM acquisition of fluorescently labelled cells or subcellular structures with micrometer precision. The strategy relies on fluorescence preservation during sample preparation and targeting based on confocal acquisition of the fluorescence signal in the resin block. Targeted trimming of the block exposes the cell of interest and laser branding of the surface after trimming creates landmarks to precisely position the FIB-SEM acquisition. Using this method, we acquired volumes of specific single cells within large tissues such as a 3D culture of mouse primary mammary gland organoids, tracheal terminal cells in *Drosophila melanogaster* larvae and ovarian follicular cells in adult *Drosophila*, discovering ultrastructural details that could not be appreciated before.

**Summary:** Ronchi et al. present a workflow to facilitate the precise targeting of three-dimensional (3D) Electron Microscopy acquisitions, guided by fluorescence. This method allows ultrastructural visualization of single cells within a millimeter-range large specimen, based on molecular identity characterized by fluorescence.

## Introduction

The blooming of new technologies and workflows over the last decade has dramatically increased the value of electron microscopy (EM) for cell biology. Volume SEMs have opened the possibility to acquire large volumes (Peddie and Collinson, 2014; Titze and Genoud, 2016). Serial Block face SEM (SBEM – Denk and Horstmann, 2004) and Focused Ion Beam SEM (FIB-SEM – Heymann et al., 2006, 2009; Knott et al., 2008; Hekking et al., 2009) offer the possibility to image volumes at nanometer resolution in a semi-automated way, complementing the tedious and highly technically demanding exercise of serial sectioning practiced in many EM laboratories. Both instruments combine iterative slicing and SEM imaging. While SBEM uses a diamond knife to remove thin sections from the block surface, in the FIB-SEM, an ion beam (most often Gallium ions) is used to ablate thin layers of material. In both cases, the electron beam is scanned onto the freshly exposed sample surface to produce images built from the signal collected by electron detectors (either backscattered or secondary electrons). The iteration of imaging and milling over thousands of cycles generates a stack of images that are then digitally aligned and combined to reconstruct a volume (Narayan and Subramaniam, 2015). For both techniques, the achievable X,Y resolution is comparable (3-4 nm). However, the use of an ion beam to remove material enables finer sectioning resolution, down to 3-4 nm (Wei et al., 2012; Xu et al., 2017; Hoffman et al., 2020; Müller et al., 2021). This range of slice thicknesses is not achievable with a diamond knife. For this reason, while SBEM is mostly suitable for the imaging of very large volumes (tissues or small organisms) in a non-isotropic fashion, FIB-SEM is currently considered the best suited option to automatically acquire relatively small volumes (ranging from subcellular to a few cells) at high isotropic resolution.

Specific sample preparation protocols have been designed for FIB-SEM and SBEM (e.g. Deerinck et al., 2010; Maco et al., 2013; Hua et al., 2015). Common to all of them is the requirement to introduce high amounts of heavy metals *en bloc* during sample processing. The electron scattering properties of heavy metals such as osmium, uranium and lead produce high image contrast. At the same time, metals ground the electron charges accumulating during electron imaging. Another important aspect of sample preparation concerns the choice of embedding media. The best milling performances with the FIB-SEM have been achieved so far with hard and rigid polymers. Therefore, epoxy resins such as Epon812, Hard Plus, Spurr’s and Durcupan (Kizilyaprak et al., 2015) are the most common choices.

When approaching a large specimen for 3D EM, restricting the acquisition to a sub-volume of interest is often necessary. When imaging a defined structure, e.g. a specific cell in a tissue, precise targeting allows the optimization of the acquisition time and the amount of data generated, therefore increasing the throughput of such experiments. To identify the region of interest (ROI), either morphological cues or fluorescence can be used, depending on the application. For morphology-based targeting, X rays have proven particularly interesting, because of the possibility to visualize micrometer-size anatomical cues of the specimen (Handschuh et al., 2013). However, when a structure or cell of interest can be labelled by fluorescent dyes or proteins, Correlative Light and Electron Microscopy (CLEM) becomes the most popular choice to guide EM acquisition or to integrate the ultrastructural information offered by EM with the molecular identity labelled in fluorescence (Mironov and Beznoussenko, 2009; Caplan et al., 2011; de Boer et al., 2015; Bykov et al., 2016). One way to classify CLEM techniques is to refer to the step in which the light microscopy (LM) is acquired in the EM sample preparation procedure. Typically, we distinguish two approaches: pre-embedding LM and post-embedding (or *in resin*) LM.

Although many CLEM workflows have been developed, utilizing fluorescent signals to target specific objects within a resin embedded specimen is still a challenging task. Indeed, most of the EM sample preparation protocols, especially those for volume SEM imaging, require both large concentrations of heavy metals and heat-polymerized epoxy resins. Both these factors affect the fluorescence: metals by quenching fluorophores in their vicinity and heat-polymerized hydrophobic epoxy resins by dehydrating and denaturing fluorescent proteins (FPs) and exhibiting non-optimal properties for LM (Paez-Segala et al., 2015). Therefore, such protocols are not compatible with fluorescence preservation. As a consequence, for FIB-SEM targeting, pre-embedding LM is often necessary. However, staining, dehydration and embedding induce anisotropic distortions to the sample (Zhang et al., 2017), and therefore the location of a structure to be acquired by EM from a LM volume cannot be predicted with sufficient precision. To mitigate this difficulty, a third imaging modality has been introduced in the workflow (e.g. X-ray micro computed tomography - μCT) to reveal the local distortion introduced during sample preparation (Karreman et al., 2016). This has proven helpful to predict the position of ROIs after sample preparation, especially for 3D targeting in brain samples. However, this complicates the workflows and requires access to additional expensive equipment. Moreover, most available benchtop X-ray machines have limited resolution, lower than achievable by LM, and not all samples offer enough morphological cues that can be recognized in the μCT scan.

To overcome the problem of sample deformation and to precisely assign the location of a fluorescently-tagged structure in an EM image, new strategies have been identified to preserve the fluorescence during sample preparation, which allow post embedding CLEM (Nixon et al., 2009; Kukulski et al., 2011; Peddie and Collinson, 2014; Biel et al., 2003). In such approaches, fluorescence preservation is enabled by reducing the amount of heavy metals in the sample, which comes with a compromise for EM contrast. Typical protocols avoid osmium and use only a small amount of uranyl acetate (UA) to stain the sample. The best suited resins to retain fluorescence are methacrylates (e.g. Lowicryl HM20 – Armbruster et al., 1982), because they are less hydrophobic than epoxy resins and can be UV-polymerized at low temperatures. This has the additional advantage of reducing the heat-induced denaturation of the fluorescent proteins. These approaches have been developed and utilized mostly for transmission electron microscopy (TEM) applications. The possibility of imaging the same field of view on the same sections at the light microscope before moving to the TEM, increases the accuracy of correlation for on-section CLEM (Kukulski et al., 2011; Avinoam et al., 2015). However, TEM imaging techniques, even when combined with CLEM, have limited power for acquiring large volumes in 3D as they would depend on the tedious serial sectioning and large scale imaging of serial sections (Mathew et al., 2020). Therefore, finding an easy workflow to combine post embedding LM and an automated volume SEM would increase the throughput of such experiments.

Recent publications showed good results in FIB-SEM acquisition of samples that were high pressure frozen, freeze substituted with low amount of UA and embedded in acrylic resins (Höhn et al., 2015; Porrati et al., 2019). Based on these observations and on our previously established sample preparation protocols for on-section CLEM of tissues (Hampoelz et al., 2016, 2019; Wong et al., 2020; Lee et al., 2020), we have established an easy and robust workflow that allows targeted FIB-SEM imaging of fluorescently labelled structures in a large volume (~7mm^2^ x 200/400 μm depth). We show that FIB-SEM targeting can be achieved with micrometer precision, based exclusively on fluorescence, with no need to rely on characteristic anatomical or morphological features. Despite the low amount of heavy metals in the sample and the lower hardness of the resin used, the imaging quality enabled fine ultrastructural analysis, proving that FIB-SEM is compatible with a large spectrum of sample preparation procedures.

## Results

### Sample preparation

To develop a sample preparation strategy that would best combine EM ultrastructure quality and fluorescence preservation for targeting, we used as a starting point the protocols we had previously optimized for on-section CLEM experiments (Hampoelz et al., 2016). We high-pressure froze the samples (mammary gland organoids in Matrigel, *Drosophila* ovaries, and dissected *Drosophila* larvae) and freeze substituted them with 0.1% UA in acetone (table 1). After a 72h long incubation at −90°C (the time may vary for different samples), the temperature was increased to allow the UA to stain the biological material. We found that an optimal concentration of UA in the sample (the best compromise between EM contrast and fluorescence preservation) was achieved by increasing the temperature to −45°C with a speed of 3°C /h and then incubating the samples in the UA solution for additional 5h at −45°C. The samples were then rinsed with pure acetone before infiltration with the resin Lowicryl HM20. As previously observed for on-section CLEM (Hampoelz et al., 2016, 2019; Lee et al., 2020; Wong et al., 2020), this sample preparation method preserved the fluorescence of the samples. Moreover, it proved suitable for ultrastructural imaging not only for TEM applications, but also for FIB-SEM for several heterogeneous samples (Fig. 1). Even though we expect that the preservation of other fluorescent proteins will be similar (especially for red and far red fluorophores), for all the experiments presented in this paper we have used genetically engineered mCherry or DsRed-tagged proteins. With this experimental setup, we could obtain fluorescence levels that were visible at several hundreds of microns depth within the resin block, when scanning with a confocal microscope over the entire block (Fig. 1a, 1g and 1k). Moreover, this sample preparation was compatible with FIB-SEM imaging, as we could achieve good imaging and milling quality for large volumes (~80×60×80 μm^3^, Fig. 1b, 1h and 1l), with sufficient contrast to visualize subcellular structures, when imaging at 8 or 10nm voxel size. For example, we were able to visualize membranebound organelles, such as ER (Fig. 1n), multivesicular bodies (Fig. 1d and 1o), mitochondria (Fig. 1e and 1p; cristae visible in fig. 1e), Golgi apparatus (Fig. 1c and 1i), but also membrane invaginations (Fig. 1m), nuclear pores (Fig. 1j), centrioles (Fig. 1f) and single microtubules (Fig. 1e).

**Figure 1.**
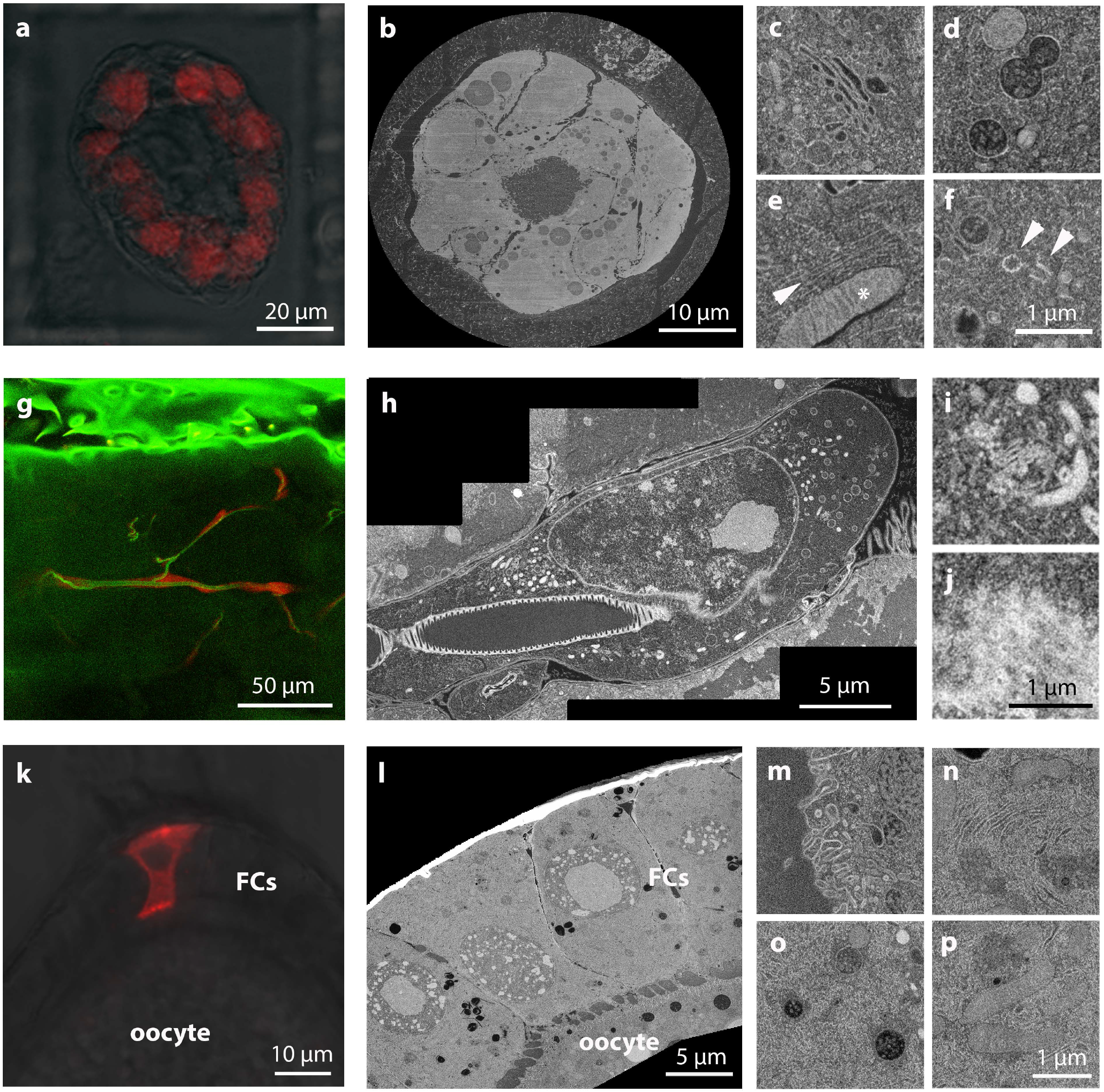
Sample preparation provides optimal fluorescence preservation and FIB-SEM imaging quality. a-f) Primary mammary gland organoids expressing H2B-mCherry (red). a) confocal image acquired from the resin block. In red the mCherry signal, overlaid to the bright field image. b) slice of the FIB-SEM volume of the entire organoid shown in a, acquired at 15nm isovoxel size. Due to the different orientation of the confocal and FIB-SEM acquisition, the view in b is orthogonal to the view in a. c-f) high magnification details of single cell volumes acquired from other organoids at 8 nm isotropic pixel size. c shows the Golgi complex; in d, multi-vesicular bodies, where we can distinguish single vesicles in the lumen; in e, a mitochondrion with visible cristae (asterisk) and a bundle of cytoskeleton filaments (probably microtubules, arrowhead); in f, a centrosome with the 2 centrioles in orthogonal orientations highlighted by arrowheads. g-j) *Drosophila* trachea terminal cell expressing cytoplasmic DsRed. a) confocal slice acquired from the resin block after sample preparation. The autofluorescence of the tissue in the green channel (including the trachea tube). In red, DsRed, specifically expressed by the trachea cells. h) slice of the FIB-SEM volume of a portion of the fluorescent cell shown in a, acquired at 10nm isotropic voxel size. I,j) details of the same volume, showing the Golgi apparatus and mitochondria (i) and nuclear pores in top view, at the nuclear envelope (j). k-p) *Drosophila* ovarian follicular cells, with clonal expression of *Dhc64C* RNAi marked by CD8-mCherry expression. a) confocal image acquired from the resin block. In red the CD8-mCherry signal, overlaid to the bright field image. Oocyte and follicular cells (FCs) are indicated. b) slice of the FIB-SEM volume of the same cell shown in a, acquired at 10nm isotropic voxel size. m-p) details of the same volume: in m, invaginations of the oocyte plasma membrane; in n, area rich in ER cisternae in a FC; in o, multi-vesicular bodies in a FC, with single vesicles in the lumen; in p mitochondria in a FC.

### Fluorescence imaging in block

After preparing the samples by high pressure freezing and freeze substitution as described above, we separated the resin blocks from the aluminum planchettes and polished the block surface by removing a few microns with a diamond trimming knife (Cryotrim 90, Diatome). This operation was done in order to remove the rings imprinted from the planchette, which could diffract the laser light, creating artefacts during the subsequent confocal acquisition. Next, the blocks were placed in a glass bottom dish (Mattek) on a drop of water. Hydration of the blocks increased the fluorescent signal close to the surface and this can be crucial for dim fluorescent samples. With this setup, we were able to acquire a tiled Z-stack confocal scan to cover the entire part of the block containing the biological sample (typically ~ 1500 x 1000 x 200/400 μm^3^), in order to identify rare targets. As fluorescence imaging in blocks has not been well characterized so far, we set out to quantitatively analyze the behavior of mCherry fluorescence in the block. For this analysis, we used mouse primary mammary gland-derived organoids expressing histone 2B (H2B)-mCherry after 10 days in culture.

Each organoid develops as a clonal expansion of a single cell. Thus, we assume that the fluorescence expression of each interphase nucleus in one organoid has similar intensity levels. First, we characterized the fluorescence intensity of H2B-mCherry in a confocal Z-stack. Fig. 2a and 2b show the example of a single representative lowicryl embedded organoid. The fluorescent signal was almost 10 times higher for nuclei exposed on the block surface, and exhibited a drastic decay in imaging planes below the surface. While there was large variability between nuclei positioned between 2 and 4 μm beneath the surface, the intensity level reached a plateau at approximately 8 μm and settled to a value of 10-15% of the intensity at the surface. Although the quantitative analysis was performed on single organoids, such that we could assume that the original fluorescence of the nuclei was homogeneous, we could detect fluorescence from the entire depth of the block, up to 400 μm from the surface, suggesting that the intensity did not drop much at further depths.

**Figure 2.**
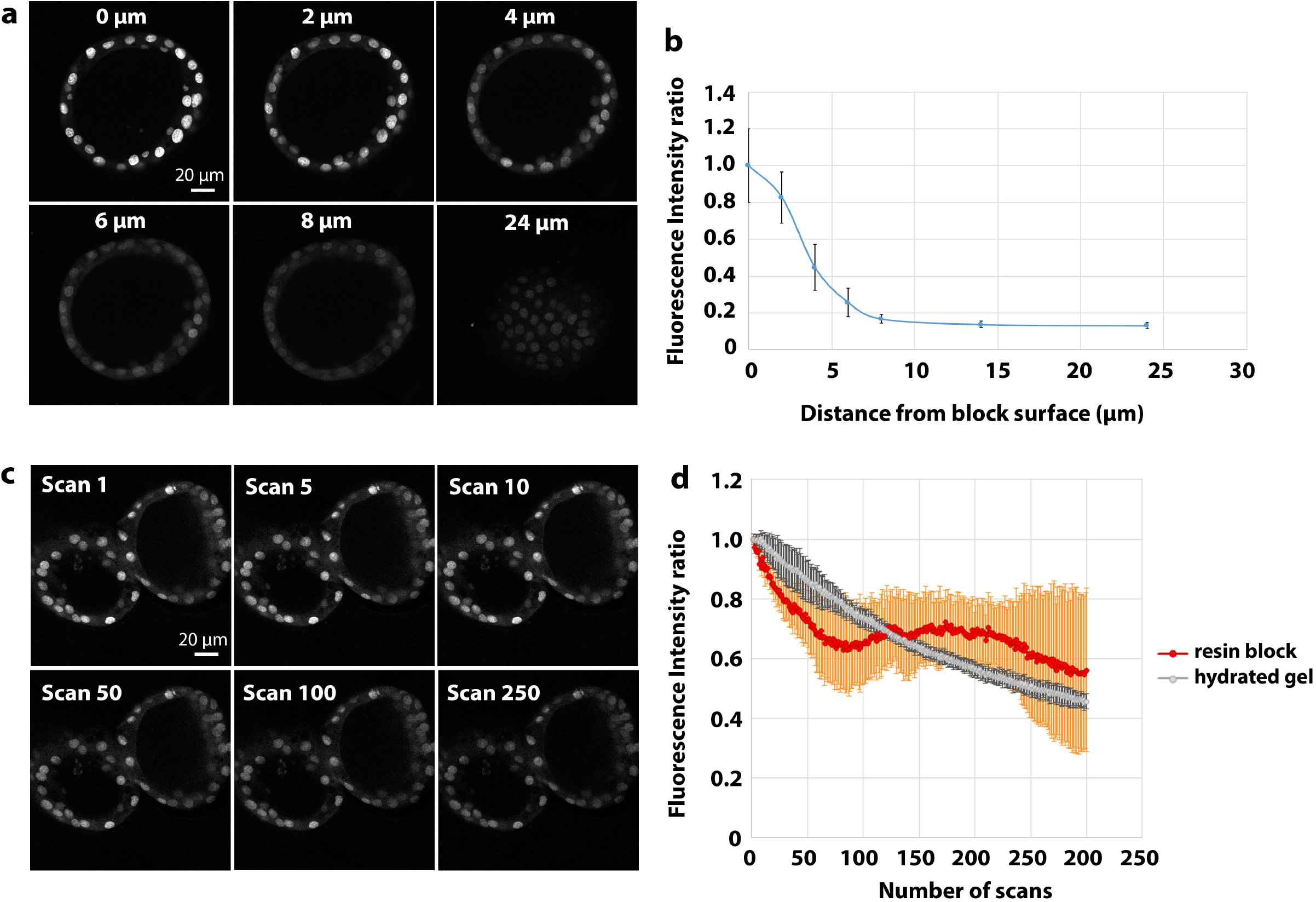
Characterization of the behavior of mCherry fluorescence in resin block. a-b) fluorescence intensity measurements of mCherry-H2B in mammary gland organoids. Confocal stack of a representative sample in a. The integrated fluorescence intensity of 10 nuclei per confocal slice was measured and the average (± standard deviation – s.d.) is plotted in b in relation to the distance from the block surface. The data are normalized to the average fluorescence intensity of the nuclei at the surface. c-d) bleaching curve of mCherry-H2B in resin block. A field of view containing organoids has been consecutively scanned 250 times, with the settings described in the methods section. Images are shown in c, and quantification of the integrated fluorescence intensity of nuclei from 3 independent organoids (± s.d.) is shown in d (red curve). Each experimental point in the chart is represented as fraction of the fluorescence intensity in the first image. The bleaching experiment of mCherry-H2B in PFA-fixed hydrated Matrigel has been conducted with the same settings used for resin embedded organoids. Average of 3 independent organoids (± s.d.) is represented in grey.

Despite the reduced fluorescence detected in deep structures, we noticed that the signal was unexpectedly resistant to photobleaching during acquisition (Fig.2c). To quantify the photobleaching of H2B-mCherry in block, we repeatedly scanned confocal sections of organoids ~2-4 μm from the block surface with the same laser settings used for imaging (Fig. 2c,d) and found that after 250 sequential scans the signal dropped on average by only ~40%. This behavior was similar to that of H2B-mCherry tagged organoid nuclei in hydrated Matrigel, which was used as a control. Although a quantitative analysis was carried out only with the organoids samples, the same resistance to photobleaching was observed qualitatively also for the other samples used in this study. In summary, even though the fluorescence signal decreases with the distance from the block surface, the resistance to photobleaching allows to target structures throughout the entire high pressure frozen sample volume, using relatively high photon doses.

### Targeting strategy and FIB-SEM imaging

FIB-SEM imaging requires precise Z targeting, since ion milling quality degrades when moving deeper from the surface of the block. Therefore, it is beneficial to have the target as close as possible to the block surface. To achieve this when the target is initially positioned deep inside the block, a good prediction of the location of the ROI and cautious trimming at the microtome are required. Our targeting approach relies on cycles of confocal scans of the block and measurements of the distance of the target from the surface, followed by removal of the resin above the target with a trimming diamond knife (Fig.3 and Fig. 4). The first confocal acquisition was performed with a low Z resolution (10μm pixel size) on the entire volume (~1000×1500×200/400 μm^3^ – Fig. 3, 4a, 4i, S1, S2a and S3a). After identification of the target, the first trimming step was performed. In view of the low Z resolution of the fluorescence acquisition, we did not remove the entire predicted excess resin thickness, but left a considerable buffer (typically, 30-40 μm). Subsequent imaging with progressively better Z resolution and trimming cycles allowed to get increasingly closer to the target, until it was a few micrometers underneath the block surface. We normally executed this approach in 3 steps, which brought the target within 30-40, then 10-15 and finally 2-5 μm from the surface, respectively (Fig. 3 and 4c). However, more steps could help in proceeding with higher confidence. While bringing the fluorescence target closer to the surface, the imaging quality improved, allowing to identify small targets with better precision (for example, single mitotic cells in an organoid - Fig. 4i-n).

**Figure 3.**
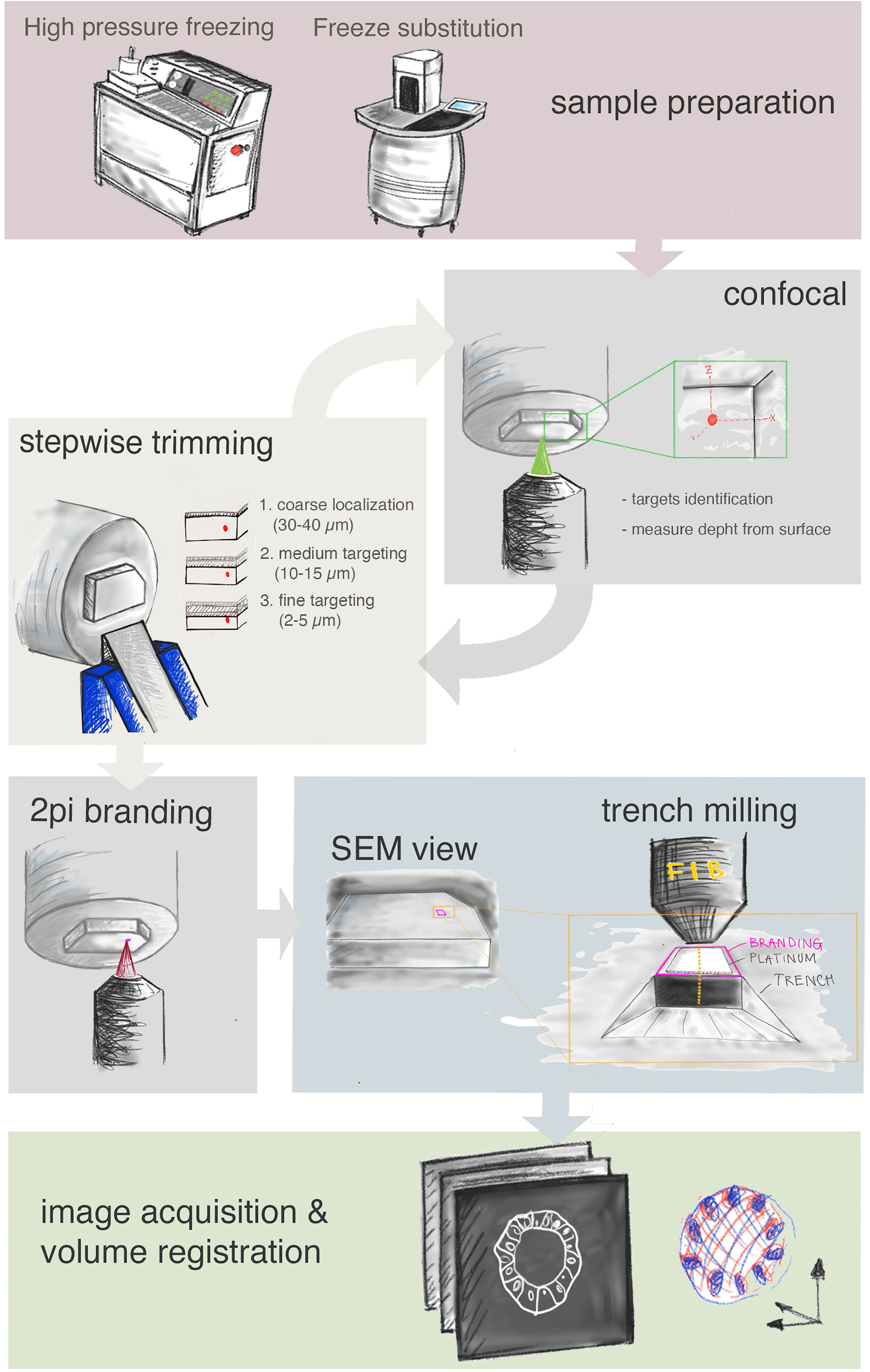
Workflow of the method. Schematic representation of the method’s main steps. From the top: sample preparation consists of high pressure freezing and freeze substitution. Second row: cycles of confocal acquisition and trimming to progressively assess the depth of the target relative to the block surface. Normally, 3 iterations were sufficient, reaching each time the approximate distance in z from target as indicated in the figure. Third row: the block surface is marked by 2 photon branding (magenta). Preparation for FIB-SEM consists of placing the platinum coating and trench milling with the ion beam in the vicinity of the branded mark. Finally, FIB-SEM imaging and image processing in the last row, include registration of the volumes obtained by FIB-SEM and confocal acquisition.

**Figure 4.**
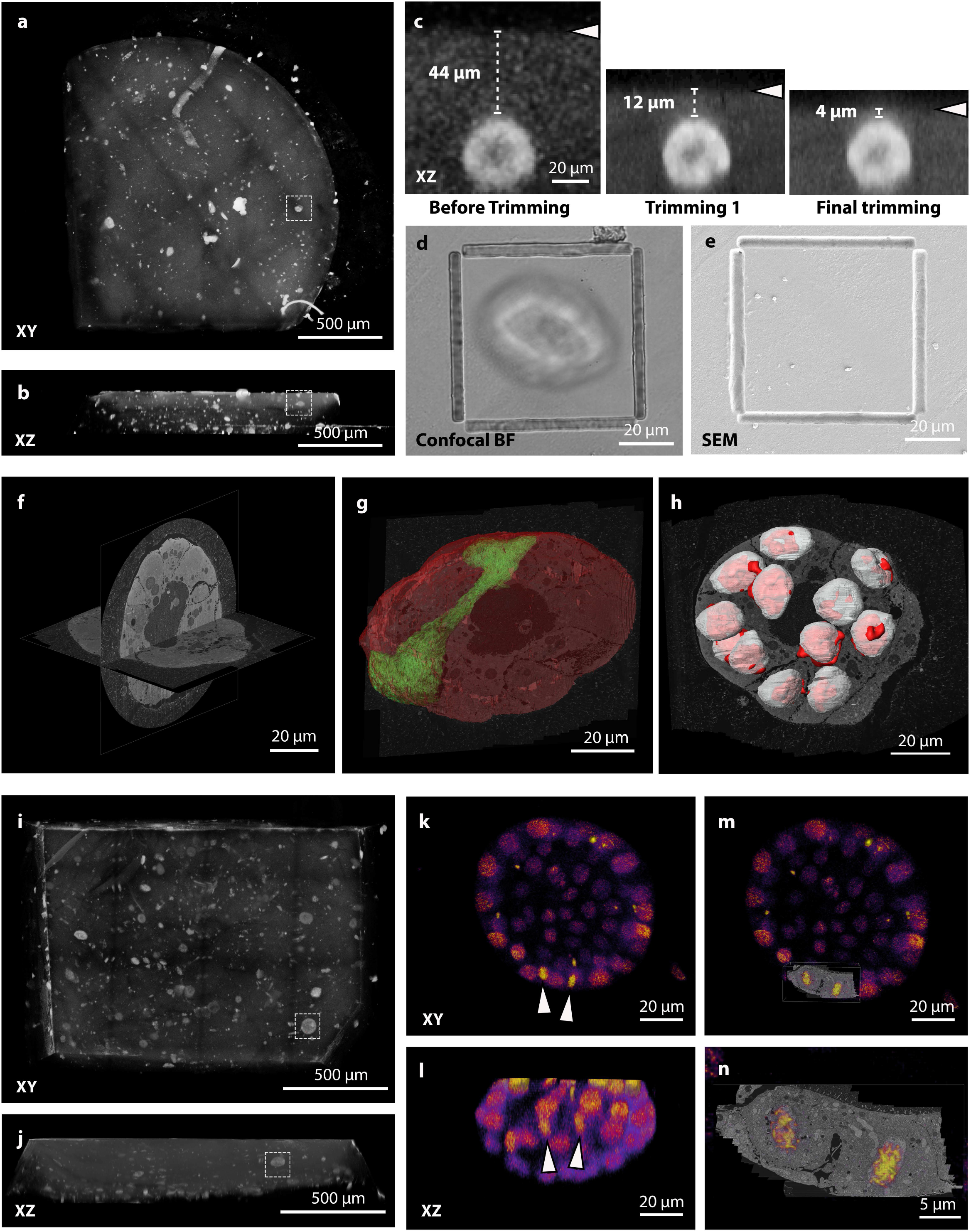
Targeting of mCherry-H2B expressing mammary gland organoids grown in Matrigel. a,b) Tiled Z-stack confocal acquisition of the resin block. The greyscale image shows a rendering of the mCherry signal. Several organoids are visible. The autofluorescence allows the identification of the block edges and surface. XY and XZ projection views of the volume are shown in a and b, respectively. The dashed box indicates the organoid of interest, which is to be targeted for FIB-SEM acquisition. c) Iterative imaging/trimming cycles to approach the region of interest (ROI) in Z. The images show XZ views of the organoid of interest and its position with respect to the block surface (arrowheads) as observed in high resolution acquisitions around the ROI. Before trimming, the target was located 44 μm from the block surface. After trimming 1, the ROI was brought within 12 μm from the surface and with the final trimming step, the target was right under the surface (4 μm). d) confocal transmitted light image of the block surface after laser branding. e) SEM image of the block surface showing the surface laser mark. Note that the biological sample is not yet exposed at the surface, making the branding the only reference to target FIB-SEM acquisition. f) FIB-SEM acquisition of the full organoid, achieved at 15 x 15 x 20 nm^3^ voxel size. Orthogonal slices through the volume are shown. g) Segmentation of the organoid (in red) and of a representative single cell (green). The cells are not distributed as a simple epithelium surrounding the lumen, but acquire a complex organization. The cell highlighted has contact to the Matrigel on 2 sides and forms the lumen with a lateral portion of its protrusion. h) Overlay of the nuclei segmented from the EM volume (white) and from the fluorescence stack (red) show precise alignment of the 2 dataset, which allows single cell identification. i-n) targeting of a mitotic telophase event within an organoid. I and j show the targeting of an organoid, as shown in a and b. After exposing the organoid, a mitotic event could be identified (arrowheads in k,l) and these cells were then targeted for FIB-SEM acquisition at high resolution (m). n shows accurate overlay between the fluorescence dataset and the nuclei of the EM volume.

After the final trimming step, the target structure is located at an optimal depth, starting 2-5 μm from the block surface. However, the block surface is millimeters in size and X,Y targeting needs to be accurate as well. Indeed, milling and imaging of large volumes with a FIB-SEM is very time consuming and long acquisitions often result in imaging instability as well as in the need for gallium source heating and consequent loss of quality in volume imaging. Therefore, a trench as small and precise as possible in X,Y is desirable. In the absence of landmarks at the surface of the trimmed block, as was the case for the organoid culture, positioning the target volume in X and Y can only be done using the distant block edges as a reference, which can be rather inaccurate. To facilitate precise X,Y targeting, we implemented a universal workflow by branding the surface of the block by two-photon (2Pi) laser following the confocal imaging. This results in an embossed feature that can be easily identified at the SEM. We therefore used this as a landmark to define the positioning of the FIB-SEM acquisition window (Fig. 3, 4d and 4e). Although the orientation of the region to be acquired can be easily deduced using the position of the branding on the block surface, using asymmetric shapes can be useful (see for instance Fig. S1 and S2). We tried several wavelengths, laser powers and repetition numbers to this aim. The best experimentally determined setup with our microscope is described in material and methods.

The targeting of a volume of interest, including imaging, finding the target, trimming and branding, was typically achieved in 3-5 hours.

At this point, the volume of interest lies close to the surface of the block and is highlighted by the laser marks. It is therefore straightforward to acquire targeted FIB-SEM data. FIB-SEM imaging of Lowicryl embedded specimen has rarely been described, but showed in our hands satisfying results while using standard imaging parameters (see Material and methods for details) and special attention in keeping a stable field of view (FOV). Similar to the behavior of other resins used in FIB-SEM, Lowicryl proved sensitive to electron beam exposure, and changes in the FOV dimension lead to milling instabilities (i.e. curtaining artefacts) that were resolved only after a few sections. However, using the confocal volume to predict the exact location of the FOV to be acquired, we were able to define an acquisition window that remained stable throughout the acquisition. We noticed that the images showed adequate contrast despite the minimal amount of heavy metals in the sample. We were able to visualize membrane-bound organelles and other subcellular structures when we acquired single (or few) cells at 8-10nm isotropic pixel resolution. A compromise in resolution had to be made for very large FOV, in order to reduce the acquisition time and to avoid milling instabilities due to an excessive electron dose. For instance, acquisition of full organoids was possible at 15×15×15/20nm^3^ voxel size.

This method therefore proved to be suitable for a large spectrum of applications, ranging from large volumes (e.g. entire mouse primary mammary gland organoids, which are spheroids up to 70 μm in diameter; Fig. 4a-h) to single cells (mitotic telophase in an organoid within a whole 3D cell culture – Fig. 4i-n - or single follicle cells in *Drosophila* oocytes – Fig. S3).

Recent work showed that in resin fluorescence can be recovered after SEM acquisition if sections are re-hydrated (Peddie et al., 2017). We therefore tested if we could target a second structure after the first one had been imaged by FIB-SEM (Fig. S1). For this experiment, we used a block containing *Drosophila* ovaries. Simultaneous expression of mCherry and dynein heavy chain siRNA (small interfering RNA) was induced in sparse follicle cell clones in a mosaic fashion. After a confocal scan of the block, we could identify 2 groups of cells, located at different depths from the surface (Fig. S1c). After targeting and acquiring the cluster closer to the surface as described above, we aimed for the second group. We thus removed the part of the block containing the first acquired volume using a razor blade (Fig. S1g). This was necessary to avoid damaging of the trimming knife, due to the hardening of the milled resin. Confocal imaging of the remaining part of the block confirmed that the second target was indeed still visible and comparable in brightness to its level before FIB-SEM of the first target (Fig. S1j). We therefore approached this second group of cells according to our workflow. This experiment shows that several structures of interest, also such that are located at different depth, within a single block can be sequentially targeted and acquired by confocal microscopy and FIB-SEM.

### Acquisition of *Drosophila* larval tracheal cells

Having developed and characterized the method, we applied it to investigate biological samples that would be otherwise difficult to approach by 3D EM. The first problem we addressed was the characterization of the terminal cells of the tracheal system in *Drosophila* larvae. These cells form branches on the surface of oxygen-demanding tissues such as muscles. An intriguing aspect is the cells’ striking organization of membrane domains. As epithelial cells, their basal membrane faces the target tissue and forms the outside of the branches, while their apical membrane is invaginated to form an intracellular tube. The cells are supported by a collagen-containing extracellular matrix (ECM) on the basal side and a chitin-containing ECM on the apical plasma membrane (aECM) that resembles the epidermal cuticle. The tracheal aECM forms ridges known as taenidia, which line the perimeter of the tubes and confer the physical rigidity that prevents the tubes from collapsing. The ability of tracheal cells to secrete this aECM is thus crucial to the proper development and survival of the animal (Öztürk-Çolak et al., 2016a). An ultrastructural characterization that is necessary to understand the physiology and development of tracheal terminal cells can only be obtained using EM. Previous studies were limited to 2D TEM at embryonic development, or the earliest larval stages (Itakura et al., 2018; Öztürk-Çolak et al., 2016b; Jones et al., 2014; Nikolova and Metzstein, 2015) and the ultrastructure of the tracheal ECM has thus never been observed at the latest larval stage, even though this is when the terminal cells undertake most of their growth (JayaNandanan et al., 2014). Due to the size of the animal and the small number of tracheal terminal cells, a 3D EM analysis of these cells at later stages requires a precise targeting strategy (Fig. S2a,b), which was so far difficult to accomplish. On the other hand, these cells have a complex 3D organization, which can be best appreciated with a 3D EM method. We therefore studied them with FIB-SEM and used the method described above to target them.

We used flies expressing DsRed in all tracheal cells using the Gal4/UAS transgenic expression system. The larvae were dissected and prepared as described in Material and Methods. The fluorescence was preserved in all cases (Fig. 1 and Fig. S2) and we identified several ROIs in each sample, where the nuclei of the cells and/or branching points of the trachea tube were visible (Fig. S2, Fig. 5). The volumes of interest ranged from few microns from the surface to up to 80μm (which required precise Z targeting by confocal imaging and trimming – Fig. S2c,d). After trimming, we used the 2Pi branding (Fig. S2e,f) to easily retrieve the location of the region of interest (ROI) in X,Y on the block surface in the SEM. We then performed the FIB-SEM acquisition of the ROIs (Fig. S2f,g) and in all cases we acquired with high precision and high confidence the targeted structures (Fig. 5a,b). The 3D analysis of such areas revealed the 3D organization of the aECM in the tracheal tube, showing for the first time its organization at the branching points (Fig. 5a,c,d) and a novel organization of the taenidial ridges (Fig. 5c,d). While in all previously published samples analyzed by TEM (embryos and first-instar larvae), the taenidia appear as knobs (Öztürk-Çolak et al., 2016b; Itakura et al., 2018) or ridges (Nikolova and Metzstein, 2015) in their cross-section, in our data the shape in the third larval stage is reminiscent of teeth (Fig. 5c,d,e). The aECM of larger multicellular branches showed the same tooth-like structures (data not shown). This was consistent across all samples (n=8), suggesting that the morphology of the tracheal aECM of third-instar larvae differs from that of the earlier developmental stages.

**Figure 5.**
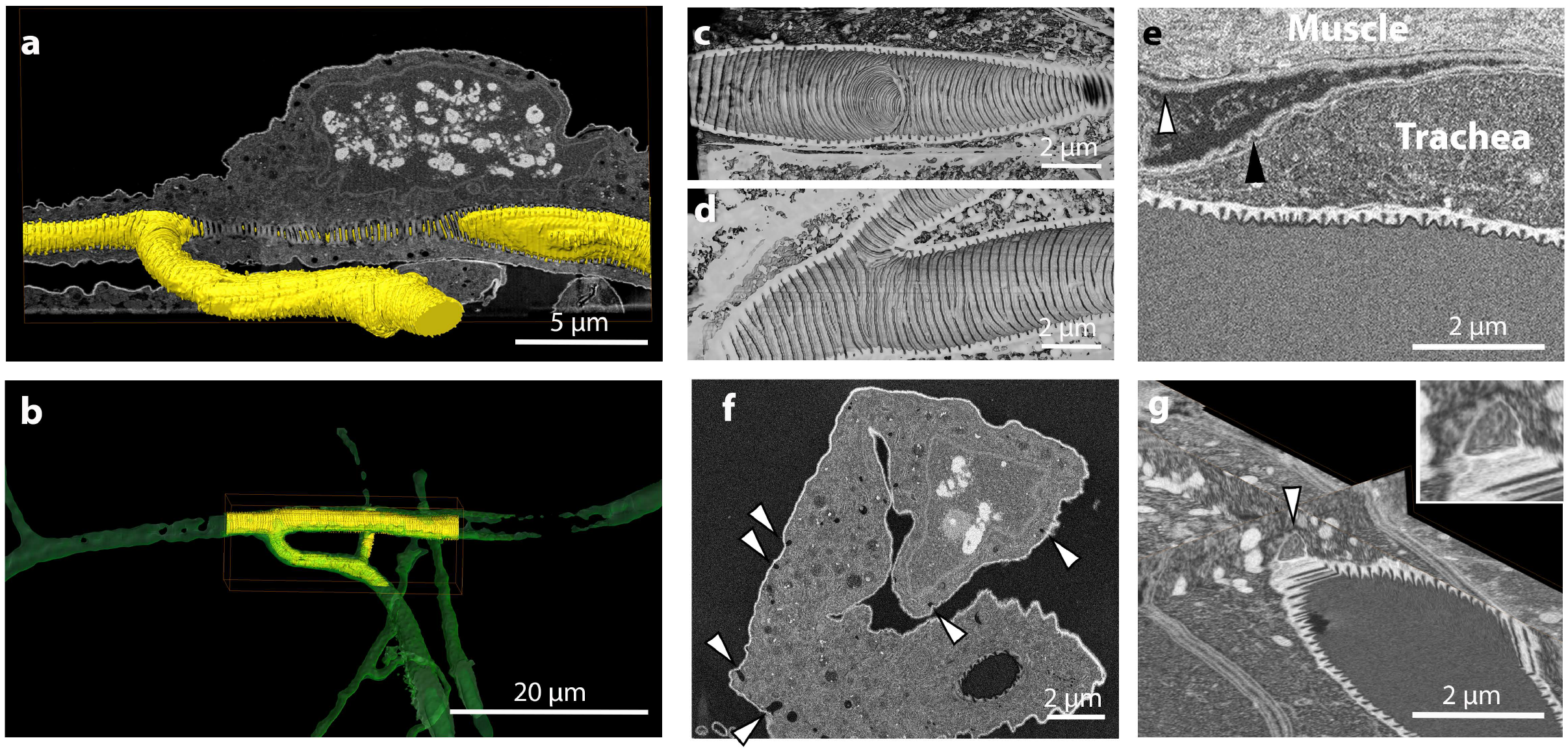
Imaging of the terminal cell of the trachea in *Drosophila* larva. The FIB-SEM acquisition of tracheal terminal cells is done as shown in Fig. S2. Here, we show representative images from different cells, to highlight interesting biological findings. a) segmentation of the hollow space inside the trachea tube (yellow) visualized with a raw image of the FIB-SEM acquisition (greyscale). b) Overlay of the segmentation of the trachea tube obtained from the confocal dataset (green) and from the FIB-SEM data (yellow). The perfect overlap confirms the accuracy of the targeting. c,d) volume rendering of the inside of the trachea tube, showing the ECM structures formed in sites of tube branching. e) FIB-SEM image, showing the organization of the basal lamina surrounding the muscle cells (white arrowhead) and the tracheal cell (black arrowhead). f) FIB-SEM image showing a cross section of a tracheal cell. Arrowheads point at invaginations of the basal plasma membrane reminiscent of endocytic activity. g) 3D visualization of a site of fusion of a carrier vesicle containing electron-scattering structures (arrowhead) with the apical plasma membrane of a trachea cell.

We observed that tracheal and muscle cells each have their own basal lamina and that the contact surface therefore shows a double layer of these basal ECM sheets separating the plasma membranes (Fig. 5e), unlike what happens in the wing disc, where tracheal branches are found encapsulated within the target tissue’s basal lamina (Guha et al., 2009). Gas exchange must thus take place through both layers of ECM, as well as through the varying thickness of the tracheal cell cytoplasm. Moreover, when we characterized the cells more closely, we also observed hallmarks of cells undergoing active endocytosis on the basal side (Fig. 5f). While it is known that tracheal cells receive proteins to secrete into the apical tube from other organs, and thus must take them up on their basal membrane first (Dong et al., 2014), this was only shown for embryonic development. It is unknown what function basal endocytosis might serve at this late larval stage. Moreover, we occasionally observed structures that might represent fusion events of carrier vesicles containing electron-scattering material with the apical plasma membrane (Fig. 5g), consistent with, for example, the delivery of material that constitutes the ECM of the trachea tube.

### Acquisition of *Drosophila* ovarian follicular cells

The *Drosophila* follicular epithelium (FE) is a monolayer of somatic epithelial cells that encapsulate the developing germline cyst and induces polarization and growth of the oocyte during oogenesis. During mid and late oogenesis, the follicle cells (FCs) that surround the oocyte differentiate into a secretory epithelium with an apical domain proximal to the oocyte, and are responsible for the apical secretion of a subset of yolk proteins and all three eggshell components (vitelline membrane, wax layer, chorion). Thanks to the possibility of generating genetic mosaics, FCs are a particularly good system to study the effect of deleterious mutations that otherwise would disrupt embryonic development. Among the variety of methods to generate genetic mosaics, the FLP-out technique allows the permanent expression of a gene of interest (e.g. Gal4) in a small subset of cells upon a short heat-shock treatment (Struhl and Basler, 1993; Pignoni and Zipursky, 1997). Once expressed, Gal4 in turn activates expression of any UAS-regulated transgene, including fluorescent markers (mCherry, GFP). In this way, the FLP-out system allows generation of fluorescently-marked, mutant FC clones adjacent to unmarked, wild-type cells, allowing direct phenotypic comparison. Although light microscopy studies of such cells are easy and informative for certain phenotypes (e.g. polarity defects - Goode and Perrimon, 1997; Bilder et al., 2000; Lu and Bilder, 2005), ultrastructural analysis of mutant clones requires a precise targeting to be able to distinguish the cells of interest from the neighboring ones in the epithelium. We therefore applied our targeting method to acquire FIB-SEM volumes containing mCherry-expressing clones together with a few adjacent wild-type, unmarked cells (control cells).

To this aim, we generated FC follicle cell clones marked by the expression of CD8-mCherry in which cytoplasmic dynein (*Dhc64C*) was knocked down by genetic RNAi (see Materials and Methods) and acquired them by FIB-SEM to assess the cellular phenotype caused by lack of dynein. Cytoplasmic dynein is essential in *Drosophila* epithelia for apical RNA localization, establishment/maintenance of apical-basal polarity, and biogenesis of microvilli by the apical targeting of Cadherin 99C (Wilkie and Davis, 2001; Swan et al., 1999; Horne-Badovinac and Bilder, 2008; D’Alterio et al., 2005; Schlichting et al., 2006). *Dhc64CRNAi* clones marked by CD8-mCherry expression in stage 10 egg chambers displayed all previously described defects associated with the lack of cytoplasmic dynein, including reduced microvillar length and aberrant funnel-like cell shape (see Fig. 6h). Interestingly, we found that the decreased length of microvilli was associated with reduced deposition of electron-scattering material such as vitelline material apically, which fails to coalesce and form an even layer of vitelline membrane as seen in the intercellular space that lies apical to wild-type FCs (Fig. 6a,d,e). Although the presence of an uneven layer of vitelline membrane was previously described in *Cad99C* mutants (Schlichting et al., 2006), it was linked to a failure of microvilli to correctly coalesce vitelline bodies into vitelline membrane. However, we noticed that *Dhc64CRNAi* cells also display an accumulation of electron-scattering material, resembling vitelline material, in the basolateral extracellular space (Fig. 6a-c). Taken together, these results suggest that the defects observed in the apical membrane of *Dhc64CRNAi* cells might result from a combination of aberrant formation of microvilli and mistargeting of vitelline material to the basolateral domain. Cytoplasmic dynein has also been described to regulate trafficking of the endo-lysosomal system (Reck-Peterson et al., 2018). In several cell types, dynein regulates microtubule minus end trafficking of late endosomes/lysosomes in coordination with the kinesin motor protein, which directs microtubule plus end transport of the vesicles (Cabukusta and Neefjes, 2018). We noticed that in *Dhc64CRNAi* cells the apically-localizing

**Figure 6.**
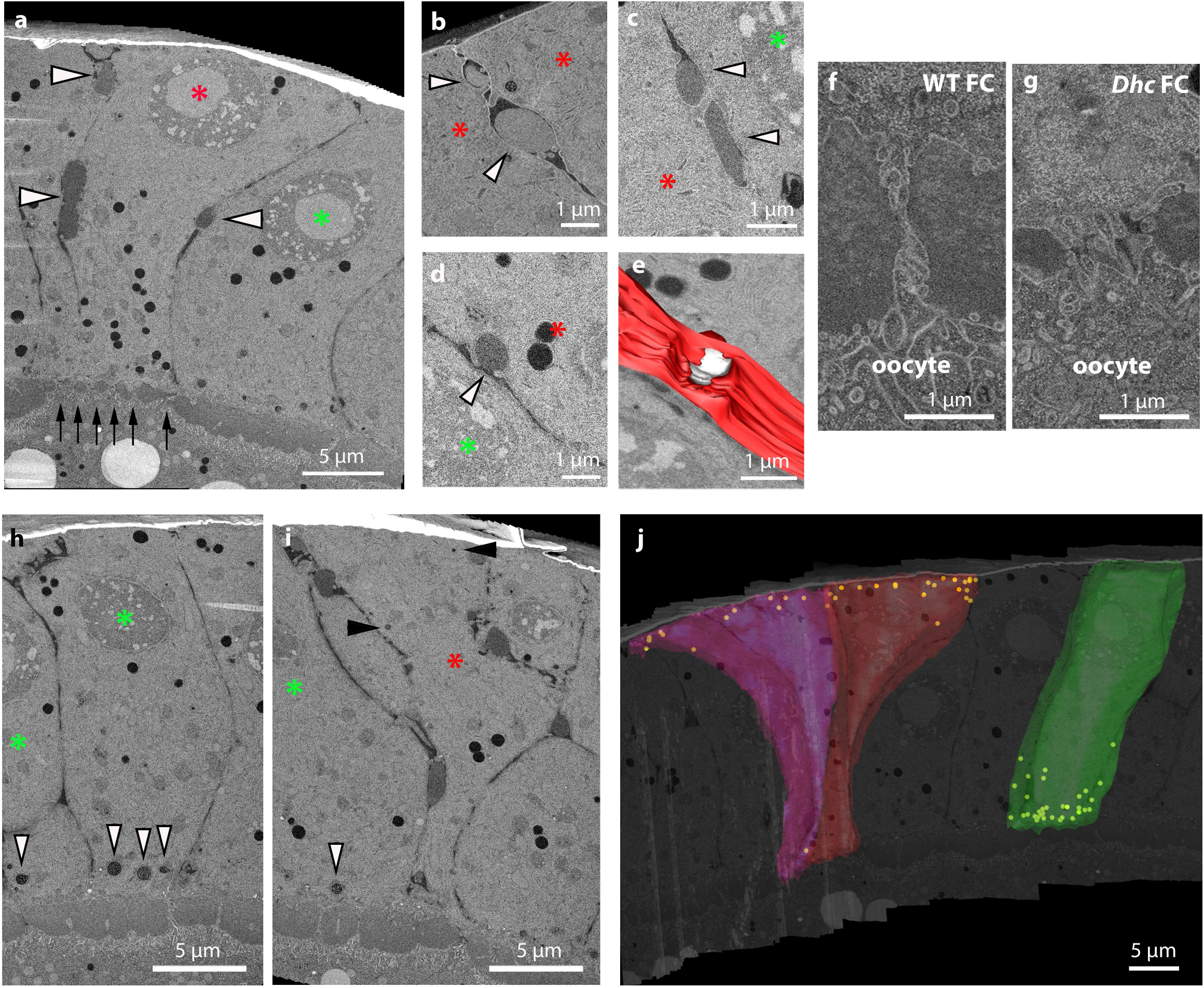
Imaging of *Dhc* KD cells in the follicular epithelium of *Drosophila* ovaries. The targeting and acquisition was done as described in Fig. S3. Identification of the Knock Down (KD, red asterisks) and WT cells (green asterisks) was done after overlay of the FIB-SEM dataset with the fluorescence stack, as shown in Fig. S3h-m. Here we show a gallery of representative images from different cells to highlight novel biological findings. a) Overview of the epithelium. Black arrows highlight the reduced space between the oocyte and the KD cell, compared to the neighboring WT epithelium. White arrowheads point to sites of lateral deposition of vitellin membrane-like electron-scattering material. b,c) higher magnification of material deposited on the lateral side between two KD cells. The material forms an electron-scattering drop (arrowheads) and its density may vary, but it does not mix with the surrounding extracellular space, which appears darker. d) potential exocytosis event of the electron-scattering material from the lateral side of a KD cell (arrowhead). e) manual segmentation of the event in d shows its 3D appearance. Plasma membrane of a KD cell is shown in red and the drop of material secreted in white. f,g) different extension of the microvilli between follicle cells and oocyte in a WT (f) and a KD cell (g). h-j) distribution of multi vesicular bodies (MVB). A WT cell accumulates large MVBs on its apical side (h, white arrowheads), whereas in KD cells the MVBs are mostly localized towards the basal side (i, black arrowheads). j shows a 3D render of a segmentation of 2 KD cells (red and purple) and a WT cell (green), overlaid on a 2D slice from the FIB-SEM stack. In yellow are indicated the positions of all the MVBs of the segmented cells. The funnel-like shape of the KD cells is also particularly evident in this example.

MVBs/late endosomes cluster close to the basal membrane (Figure 6f-h), indicating the requirement of dynein in the apical localization of late endosomes in FCs. The localization of MVB basally, where microtubule plus ends are enriched (Bacallao et al., 1989; Clark et al., 1997), suggests that in *Dhc64CRNAi* cells MVB become basally localized by the plus end-directed kinesin motor. Further studies will be required to investigate this phenomenon.

## Discussion

In this paper, we present an easy and reliable workflow to target FIB-SEM volume imaging. We have targeted in total 3 complete H2B-mCherry labelled organoids, 2 single mitotic events within an organoid, 8 *Drosophila* tracheal terminal cells expressing cytoplasmic DsRed, 6 clones of *Drosophila* ovarian follicular cells knocked-down for dynein heavy chain expressing CD8-mCherry, with 100% success rate.

Compared to other targeting methods based on morphological and anatomical cues, this workflow has the advantage of directly using the molecular identity provided by fluorescent labelling of cells of interest. This has been possible so far only by integrating pre-embedding fluorescence and FIB-SEM volumes. However, such correlation can be very challenging due to anisotropic shrinkage of the samples during EM sample preparation. X-ray μCT can aid in the identification of such distortion, to allow indirect prediction of the location of the volume of interest, and it has been successfully used to target FIB-SEM volumes (Karreman et al., 2016). However, preserving the fluorescence in the sample and being able to image the very same volume by 3D light and 3D SEM techniques has a clear advantage in terms of precision and ease of application of these complex experiments. Moreover, with our workflow we are able to target the FIB-SEM acquisition of a cell starting from a millimeter-size EM block in ~3h, while Karreman et al. reported ~2 days for the X-ray based targeting (Karreman et al., 2016).

Given the high resolution of fluorescence imaging and the fact that the sample is not altered between the two imaging modalities, we foresee that it will be possible to target even subcellular structures. However, such application would require the development of strategies to align the 3D datasets after acquisition with high accuracy.

The main challenge of the method is the difficulty in imaging fluorescence deep in the block. Therefore, we have so far limited its application to overexpressed exogenous proteins. The decay in fluorescence detection when moving away from the surface depends on several factors. First, the resin scatters the light. This is true especially for embedded biological samples, because they are not homogeneous and after sample preparation they contain metals. To improve this aspect, higher numerical aperture lenses can be used to detect photons scattered at wider angles. However, such lenses typically have short working distances and this limits the depth of the block that can be reached. In this study we opted for a 25x lens, with a long working distance, to be able to detect fluorescence from the entire biological sample. An important factor that contributes to fluorescence is the hydration of the fluorophores. As previously shown (Peddie et al., 2017), fluorescent proteins in vacuum do not emit photons. However, when sections are exposed to water vapour, fluorescence can be detected. We speculate that this factor could explain the rapid drop of fluorescence intensity in the first 6-8 μm from the surface, to then stabilize to a plateau at larger depths (Fig. 2b). Therefore, we believe that fluorophore dehydration plays a major role in our fluorescence loss, probably more important than light scattering in our experiments. Another interesting observation is that fluorescence does not bleach much in resin. Upon illumination with the laser settings we used for high resolution imaging, we observed a good signal even after 250 consecutive scans. This behaviour is strikingly different from the major fluorophore photobleaching we experience when imaging resin sections. This is essential for this workflow, as it allows scanning the block several times during the iterative trimming/imaging cycles used to reach the correct Z position of the target.

Oxygen is a key player in photobleaching (Zheng et al., 2014) and the low abundance of this molecule in the resin block compared to a thin section, may explain this difference.

Another important point of our work is that sample preparation protocols compatible with fluorescence preservation have proven satisfactory for FIB-SEM milling and imaging. Despite the absence of osmium, we achieved good contrast only in the presence of 0.1% Uranyl acetate. The stability and milling properties of Lowicryl HM20 under the FIB were comparable to the Epoxy resins that are commonly used.

In summary, our data show a reliable workflow for targeting single cells within a large 3D volume based on their molecular signature. This method has enabled us to characterize in a short time specific single cells within a homogenous epithelium or in a complex volume. Given the common availability of the instrumentation required, we foresee that such a workflow provides EM laboratories new options to study cell and developmental biology in 3D.

## Material and methods

### Organoid culture

To establish the mouse strain line TetO-MYC/ TetO-Neu/ MMTV-rtTA/ R26-H2B-mCherry, mouse lines TetO-MYC/MMTV-rtTA (D’Cruz et al., 2001), TetO-Neu/ MMTV-rtTA (Moody et al., 2002) and R26-H2B-mCherry (Abe et al., 2011) (RIKEN, CDB0239K) were crossed into FVB background. Housing and care of all animals used in this study was performed at LAR (Laboratory Animal Resources) facility at EMBL Heidelberg according to guidelines and standards of the Federation of European Laboratory Animal Science Association (FELASA). All mice were bred and maintained in a 12 hours light / 12 hours dark cycle, with constant atmospheric conditions (23 ± 1 °C temperature; 60 ± 8 % humidity) and permanent access to food and water. For establishment of 3D organoids, mammary glands from 8 weeks virgin TetO-MYC/ TetO-Neu/ MMTV-rtTA/ R26-H2B-mCherry female mice, were dissected and digested following the published protocol (Jechlinger et al., 2009). Single cells were seeded after mixing them with a cold combination of Matrigel growth Factors Reduced (Corning, 356231), Rat Collagen I (RnD Systems, 3447-020-01) and PBS. Gels were allowed to polymerize before supplying them with Mammary Epithelial Cell Growth Medium (Promocell, c-21010 and supplement with Mammary Epithelial Cell Growth Supplement (Sciencell, 7652). Cultures were maintained for 3-10 days in culture in a humidified atmosphere with 5%CO_2_.

### *Drosophila* tracheal cells dissection

The *Drosophila* line used was reported previously (Best and Leptin, 2020) and carries a recombined btl-Gal4 (Shiga et al., 1996) and UAS-DsRed1 (BDSC 6282) element on the third chromosome, driving expression of DsRed in all tracheal cells. The culture was grown on standard cornmeal-agar medium at 25°C. Wandering third-instar larvae were gently collected from the vial wall using a brush and transferred to a droplet of 4°C Shields and Sang medium on a dissection plate. The larvae were filleted according to standard protocol, exposing the dorsal tracheal system attached to the skin, with all internal organs removed. After confirming that the tissue was still alive by observing the twitching of muscles on the skin, we removed the head and posterior end of the fillet. The remaining sample usually contained completely the five segments A1-A5 (corresponding to tracheal dorsal branch pairs 3-7). This was transferred directly to the high-pressure freezing carrier, prefilled with 20% Ficoll (PM70, Sigma) in Shields and Sang medium.

### Heat-shock treatment and *Drosophila* ovary dissection

The UAS-Gal4 FLP-out system was used to generate marked mutant clones in a wild-type background (Pignoni and Zipursky, 1997). Flies homozygous for an allele carrying a Gal4-inducible promoter (UAS) upstream of *Dhc64C* hairpin RNA (BDSC #36698) were crossed with *hsFlp; arm>f+>Gal4; UAS-CD8-mCherry* (kind gift from Juan Manuel Gomez-Elliff). The protocol described in González-Reyes and St Johnston (1998) was followed to generate follicle cell clones. Briefly, freshly eclosed females resulting from each cross were collected and mated with *w1118* males for 24 h at 25°C on food supplemented with yeast. Flies were heat-shocked for 1h in a water bath at 37°C, then kept at 25°C with males on yeast. Ovaries were dissected in PBS 39h post heat-shock and immediately high pressure frozen using 20% Ficoll in Schneider’s medium as cryoprotectant.

### EM Sample Preparation

All samples were high pressure frozen in their respective freezing media with an HPM010 (AbraFluid), using 3mm wide, 200 μm deep aluminum planchettes (Wohlwend GmbH). Freeze substitution and resin embedding were performed as described in the Results section in an automated AFS2 machine (Leica), using the FSP unit. To facilitate the infiltration of Lowicryl HM20 (Polysciences Inc), the temperature was gradually raised to −25°C while increasing the resin concentration in acetone. Finally, the samples were UV polymerized at −25°C. Details are shown in table 1.

### Confocal microscopy and laser branding

For confocal imaging, the blocks were mounted in a 35mm glass bottom culture dish (MatTek, Ashland, USA) immersed in a drop of deionized water. Fluorescence imaging of all the samples was done using an inverted Zeiss LSM 780 NLO microscope (Jena, Germany) equipped with a 25x Plan-Apochromat 25x/0.8 Imm Korr DIC multi immersion objective lens. First, we imaged the whole block using the “Tile Scan” function together with the “Z-Stack” function of ZEN-black. Second, from single targets we recorded high-resolution stacks to assess the overall morphology of the cells of interest. From the acquired Z-stacks we estimated the relative distance of the target to the surface of the resin block. Next, we trimmed the block using an ultramicrotome (Leica UC7) to remove excessive resin material on top of the cells of interest and iteratively repeated the imaging and trimming steps until the selected cell was positioned at a distance of ~1-5 μm to the resin block surface. Branding of the resin surface was done using the 2-photon Coherent Chameleon Ultra II Laser (Santa Clara, USA) of the Zeiss LSM 780 NLO microscope. To engrave a rectangular region on the resin block surface, we used the “Bleaching” function of ZEN black. Different settings were tested to find the optimal parameters for branding. We obtained the best results applying approximately 117mW at the sample plane of the 2-photon laser running at 765nm wavelength with an effective scan speed of 2,55μs over 50 iterations in a selected small rectangular region of the block surface. However, the success of branding and the energy level required were variable between samples, different areas on one resin block, and different sizes of branding regions. We therefore always started with minimal laser intensities and repeated the branding scan with increasing intensity until a clear brand appeared in the transmitted light image of the confocal.

### FIB-SEM acquisition

After confocal imaging and branding for targeting, the blocks were mounted on a SEM stub using silver conductive epoxy resin (Ted Pella). In case the block was later to be imaged again at the confocal to identify a second target, instead of the silver epoxy (which requires polymerization at 60°C), we attached the block to the stub using a mixture of super glue (Loctite) and colloidal silver liquid (Ted Pella). After mounting, the blocks were gold sputtered with a Quorum Q150R S coater.

For FIB-SEM acquisitions, we used a Zeiss CrossBeam XB540 or XB550, using the Atlas3D workflow (FIBICs). Briefly, a platinum coat (~1 μm thick) was deposited over the area inscribed in the laser branding. Autotuning marks were milled on the Pt surface and highlighted with Carbon. We milled large trenches with 30kV FIB beam acceleration voltage and 30nA current and polished the surface with 7 or 15nA currents. Precise milling during the run was achieved with currents of either 700pA or 1.5nA. For all experiments, the SEM imaging was done with an acceleration voltage of 1.5kV and current of 700pA, using a back-scattered electron detector (ESB). Pixel sizes and dwell times were different, depending on the volume that we acquired. For relatively small, high resolution volumes, we acquired at 8 nm isotropic pixel size (single cells in organoids) or at 10nm *(Drosophila* oocyte follicular cells, *Drosophila* trachea). For very large volumes (entire organoids – spheroids up to 65 μm in diameter) we acquired at 15×15×15/20 nm^3^ voxel size. Dwell times ranged between 8 and 12 μs, but were occasionally increased to 20 μs during the run to obtain single images with high signal-to-noise ratio.

### Image processing, visualization and segmentation

FIB-SEM image stacks were aligned using either the “Linear stack alignment with SIFT” plugin in Fiji or the “Alignment to median smoothed template” workflow recently described (Hennies et al., 2020). For visualization and registration of the different imaging modalities, we used Amira (version 2019.3 or 2020.1, ThermoFischer Scientific). The volumes of the confocal and FIB-SEM datasets were aligned using the transformation editor. When the groves left by the laser branding were visible in the FIB-SEM volume, they were used as landmarks for the registration. Otherwise, morphological features of the target cells were used (e.g. branching points of the tracheal tubes or characteristic shapes of organoids and *Drosophila* follicular cells).

## Acknowledgments

We thank the Electron Microscopy Core Facility and the Advanced Light Microscopy Facility at EMBL for their support. L. Cassella was supported by DFG-FOR 2333 grant EP 37/2-1 from the Deutsche Forschungsgemeinschaft (Germany) to A. Ephrussi. E. D’Imprima was supported by a fellowship from the EMBL Interdisciplinary programme under Marie Skłodowska-Curie Actions COFUND (EI4POD). J. Mahamid acknowledges funding from the EMBL and the European Research Council (ERC 3DCellPhase^-^ 760067).

## Author contributions

P. Ronchi conceptualized the study. P. Ronchi and P. Machado developed the method with the help of S. Schnorrenberg. P. Ronchi, P. Machado, E. D’Imprima, B.T. Best, L. Cassella and M. Garcia Montero performed the experiments. P. Ronchi, P. Machado and G. Mizzon performed data analysis and visualization. P. Ronchi, B.T. Best and L. Cassella wrote the manuscript. M. Jechlinger, A. Ephrussi, M. Leptin, J. Mahamid, Y. Schwab supervised the project. All the authors reviewed the manuscript.

