## Supplemental material (3 figures and 1 table) for "Fluorescence-based 3D targeting of FIB-SEM acquisition of small volumes in large samples"

### Supplementary figure legends

**Suppl. Figure 1. Targeting of multiple cells in the same block (*Dhc* KD cells in the follicular epithelium of *Drosophila* ovaries, expressing CD8-mCherry)**

a) workflow. b) Tiled Z-stack scan of the entire resin block in XY view, with the 2 targets highlighted in green and red. c) XZ view of the same block, with the 2 fluorescent targets segmented in the same colors. The block surface before trimming is indicated by the arrowhead. The red target is 28 $\mu$m deeper than the green one. Therefore, the 2 cells have to be exposed sequentially for FIB-SEM acquisition. d) Z targeting of the first group of CD8-mCherry positive cells (segmented in green). The image shows the position of the fluorescent cell (segmented) and the block surface (arrowhead) after trimming. e) confocal imaging of the block after trimming and laser branding around the first cells of interest (XY view). f) FIB-SEM acquisition of the same volume. The asterisks in e and f indicate the same cell viewed from orthogonal orientations. g-h) Tiled Z-stack scan of the same resin block after removing the area imaged by FIB-SEM. Note that the lower left corner of the block, previously containing the green target, is missing. The XZ view in h shows that the second target (red) is now 28 $\mu$m deep from the block surface (arrowhead). i) Z targeting of the second group of cells (segmented in red). The image shows the position of the fluorescent cell (segmented) and the block surface (arrowhead) after the second trimming. j) confocal imaging of the block after trimming and laser branding around the second cells of interest (XY view). k) FIB-SEM acquisition of the same volume. The asterisks in j and k indicate the same cell.

**Suppl. Figure 2. Targeting of tracheal terminal cell in *Drosophila* larva.**

a,b) Tiled confocal Z-scan covering the entire volume of the block containing tissue. The greyscale image shows a volume rendering of the thresholded autofluorescence signal (green channel) of the block (clearly visible is the autofluorescence of the epidermal cuticle) in XY (a) or XZ (b) views. The segmented cell of interest (in red the DsRed fluorescence of the cell, in green autofluorescence of the ECM of the tracheal tube) is highlighted in the boxed area to visualize its location in the block. c,d) in greyscale, XZ view of the volume rendering of the fluorescence of the block, with the cell of interest segmented. The arrowheads indicate the position of the block surface before (c) and after trimming (d). e) XY view of the same volume which shows the 2photon branding of the block surface marking the ROI to be acquired by FIB-SEM (segmented in magenta). f,g) Images of the block during FIB-SEM run setup: 50pA FIB image acquired with secondary electron detector (f) and 1.5keV 700pA SEM image acquired with ESB detector (g).

**Suppl. Figure 3. Targeting of a clone of *Drosophila* ovary follicular cells expressing *Dhc* RNAi and CD8-mCherry.**

a,b) Tiled Z-stack confocal acquisition of the resin block. The greyscale image shows a rendering of the red signal. A few mCherry expressing cell clones are visible (e.g. arrowhead) as well as the autofluorescence background of the Ficoll-containing freezing medium. The target of the FIB-SEM acquisition is highlighted by the dashed box and segmentation is shown to facilitate its visualization. a and b show XY and XZ views of the block, respectively. c,d) in greyscale, XZ view of the volume rendering of the fluorescence of the block, with the cells of interest segmented in 2 different colors. The arrowheads show the position of the block surface before (c) and after trimming (d). e) XY view of the same volume which shows the 2photon branding of the block surface limiting the ROI to be acquired by FIB-SEM (magenta segmentation). f,g) Images of the block during FIB-SEM run setup: 50pA FIB image acquired with secondary electron detector (f) and 1.5keV 700pA SEM image acquired with ESB detector. A crack at the interface between the basal membrane of the ovary and the empty resin is visible here. This happened frequently during the branding of *Drosophila* oocytes samples, but did not affect the structure of the follicular cells. h-m) overlays of the fluorescence (h and k) and segmented FIB-SEM (i,l) datasets show the precision of the registration in different orientations.

**<u>Supplementary table</u>**

**Table 1. Freeze substitution protocol (Leica AFS2 with FSP unit)**

| Step # | Tstart | Tend | Slope | Time | Solution | Operation | Agitation | UV |
|---|---|---|---|---|---|---|---|---|
| **1** | -90°C | -90°C | 0°C/h | 72h | FS cocktail | stay | | |
| **2** | -90°C | -45°C | 3°C/h | 15h | FS cocktail | stay | | |
| **3** | -45°C | -45°C | 0°C/h | 5h | FS cocktail | stay | | |
| **4** | -45°C | -45°C | 0°C/h | 10min | acetone | exchange/fill | | |
| **5** | -45°C | -45°C | 0°C/h | 10min | acetone | exchange/fill | | |
| **6** | -45°C | -45°C | 0°C/h | 10min | acetone | exchange/fill | | |
| **7** | -45°C | -45°C | 0°C/h | 6h | Lowicryl 10% | mix | on | |
| **8** | -45°C | -45°C | 0°C/h | 6h | Lowicryl 25% | mix | on | |
| **9** | -45°C | -35°C | 1.67°C/h | 6h | Lowicryl 50% | mix | on | |
| **10** | -35°C | -25°C | 1.67°C/h | 6h | Lowicryl 75% | mix | on | |
| **11** | -25°C | -25°C | 0°C/h | 10h | Lowicryl 100% | exchange/fill | | |
| **12** | -25°C | -25°C | 0°C/h | 10h | Lowicryl 100% | exchange/fill | | |
| **13** | -25°C | -25°C | 0°C/h | 10h | Lowicryl 100% | exchange/fill | | |
| **14** | -25°C | -25°C | 0°C/h | 48h | Lowicryl 100% | stay | | on |
| **15** | -25°C | +20°C | 5°C/h | 9h | Lowicryl 100% | stay | | on |

**Supplementary figure 1**

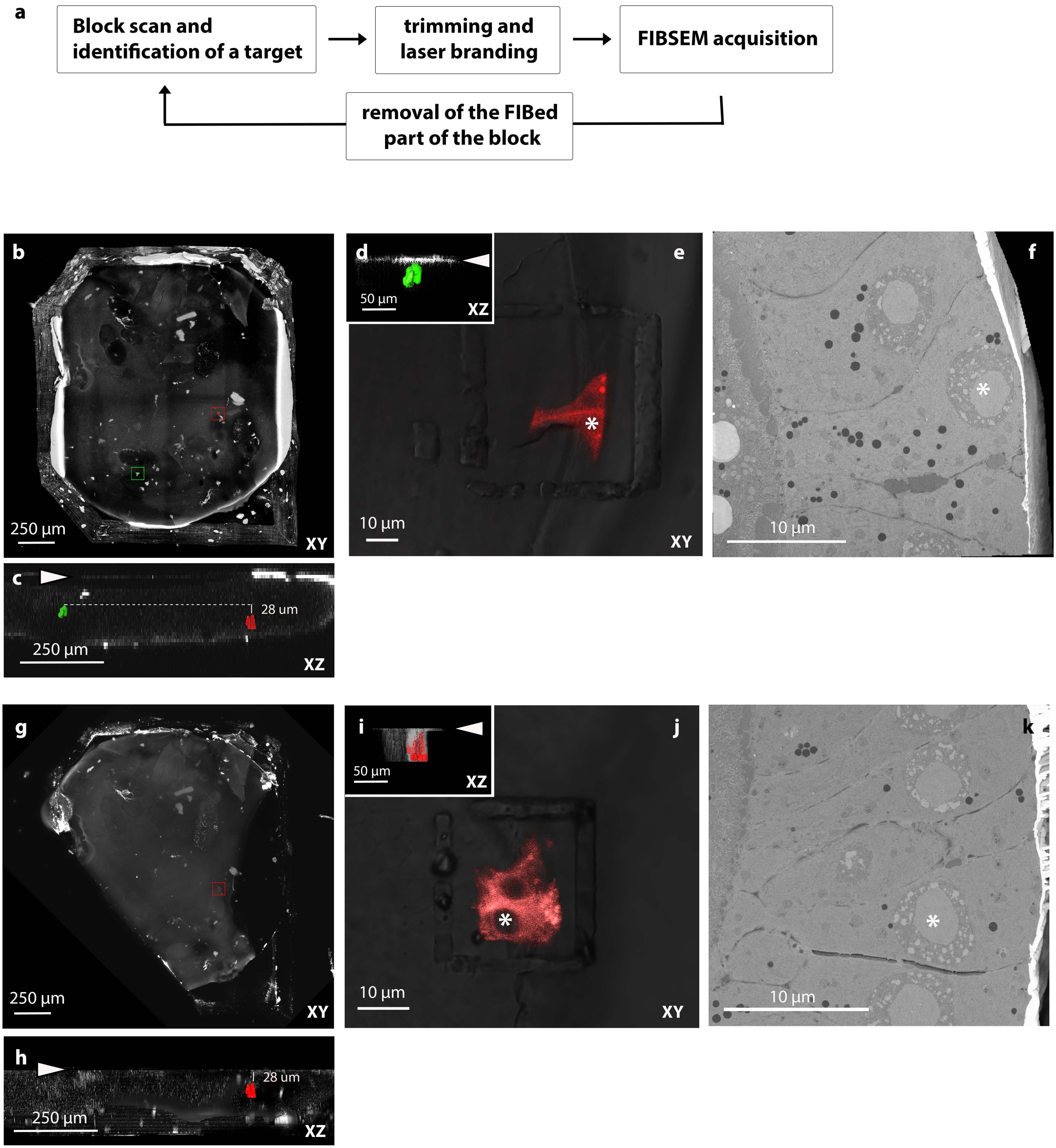

### Supplementary figure 2

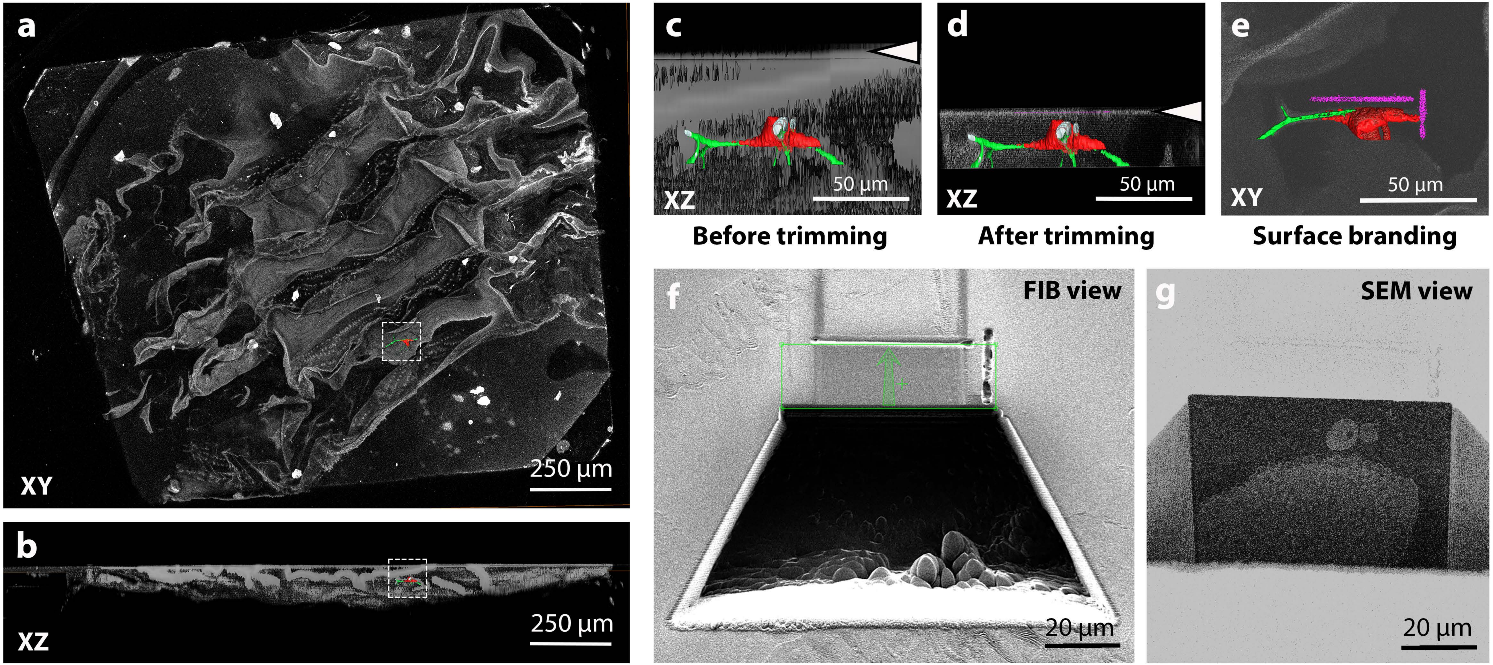

**Supplementary figure 3**

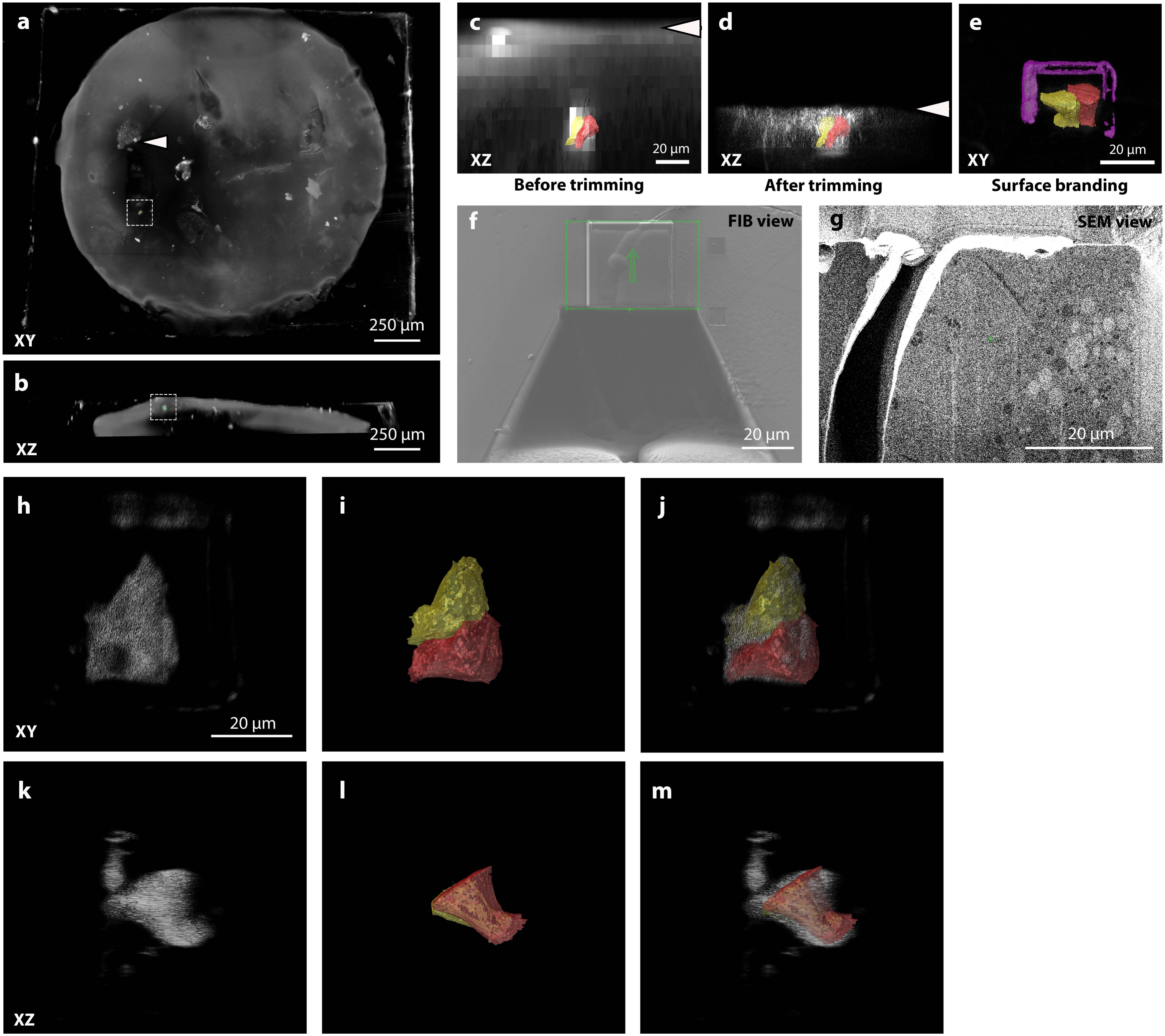
